# Genome-wide identification of *SINA* gene family in sugarcane and functional analysis of *SsSINA1a* in drought response

**DOI:** 10.1101/2022.05.01.490191

**Authors:** Jinxu Zhang, Xialan Jiang, Shenghua Xiao, Shuo Jiang, Wei Yao, Muqing Zhang

## Abstract

Sugarcane (*Saccharum* spp. hybrid) is a crucial sugar and energy crop that provides majority of the raw material for sugar and ethanol production globally. Drought represents one of the most critical constraints of sugarcane production in the subtropical parts of China. SEVEN IN ABSENTIA (SINA) act as an important E3 ubiquitin ligase and play a significant role in plant stress responses. However, the characteristics of the sugarcane *SINA* gene family have not been previously studied currently. Here, we identified 15 *SsSINA* in *Saccharum spontaneum*, 5 *ShSINA* in *Saccharum spp*. hybrid and 6 *SbSINA* in *Sorghum bicolor* based on their conserved N□terminal RING and C-terminal SINA domains, and these genes were distributed into three phylogenetic groups (I, □ and □). Collinearity analysis showed a close genetic relationship between the *SINA* genes of *S. spontaneum* and *S. bicolor*. The cis-regulatory elements in the promoter regions of the *SINA* genes were involved in a variety of plant physiological responses. Further, we identified a *SINA* gene *SsSINA1a* that significantly induced by drought stress. Overexpression of *SsSINA1a* enhanced drought tolerance in Arabidopsis through reducing leaf water loss rate. These finding indicate that SsSINA1a mediates plant drought tolerance and this study provides a new potential candidate gene for sugarcane drought-resistant breeding.

## 1. Introduction

Protein degradation mediated by the ubiquitin-proteasome system (UPS) is an important posttranslational modification, which mediated a series of cellular processes through removing misfolded and damaged proteins (Harper and Schulman, 2006; Ravid and Hochstrasser, 2008). In the UPS pathway, an ATP dependent E1 (ubiquitin activating) - E2 (ubiquitin conjugating) - E3 (ubiquitin ligase) enzyme conjugation cascade covalently links free ubiquitin moieties to target protein (Sadanandom et al. 2012). The continuously adding of ubiquitin moieties leads to the creation of a lysine-48 (K48)-linked polyubiquitinated substrate protein and subsequently recognized and degraded by the 26S proteasome (Harper and Schulman, 2006). Comparing to the relatively conserved E1s and E2s, the E3 ligases possess highly diversity and lead to diverse combination of E1s, E2s and E3s to determine the specificity of substrate ubiquitylation and strictly control countless cellular processes (Harper and Schulman, 2006). By far, at least 1000 kinds of E3 ligases in plants were identified and divided into HECT-type, RING-type and U-box-type three main classes, which widely involved in plant development and abiotic stress responses (Chen and Hellmann, 2013; Shu and Yang, 2017).

SEVEN IN ABSENTIA (SINA) belongs to a group of RING–type E3 ubiquitin ligase, which contains a cysteine-rich C3HC4-type RING finger domain in N-terminal required for mediating proteolysis of target proteins and a SINA domain in C-terminal responsible for oligomerization and substrate-binding (Hu and Fearon, 1999). The first *SINA* gene was identified in *Drosophila melanogaster* and responsible for the development of the R7 photoreceptor cell in the Drosophila eye (Carthew and Rubin, 1990; Li et al., 1997). It is generally thought that the protein sequences of SINA and SINA homologs (SIAH) are highly conserved from mammal to plant (House et al., 2003). In plants, 5 *AtSINAs* in *A. thaliana* (Wang et al., 2008), 6 *OsSINAs* in *O. sativa* (Wang et al., 2008), 6 *ZmSINAs* in *Zea mays* (Wang et al., 2008), 6 *SlSINAs* in *Solanum lycopersicum* (Wang et al., 2018), 10 *PtSINAs* in *Populus trichocarpa* (Wang et al., 2008), 11 *MdSINAs* in *Malus domestica* (Li et al., 2020) and 6 *MtSINAs* in *Medicago truncatula* (Herder et al., 2008) have been identified. In Arabidopsis, *SINA* of *Arabidopsis thaliana* 5 (*SINAT5*) negatively regulates the numbers of lateral roots by mediating the degradation of the transcriptional activator NAC1 to attenuate the auxin signal (Xie et al., 2002). *SINAT5* also participates in the regulation of florescence by mediating the ubiquitination and degradation of protein involved in flowering pathway (Park et al., 2007; Park et al., 2010). Recent years, several members from *SINA* gene family were reported to regulate plant response to drought stress (Zhang et al., 2019). The rice *SINA* gene *OsDIS1* promotes the degradation of the Ser/Thr protein kinase *OsNek6* to negatively regulate plant drought tolerance (Ning et al., 2011). *SINAT2* improves drought tolerance by activating abscisic acid (ABA)-related drought stress signals in Arabidopsis (Bao et al., 2014). However, there is few information available about the function study and the expression pattern of the *SINA* genes in sugarcane.

Sugarcane (*Saccharum* spp.) is an important crop for sugar and bioenergy production in the world (Garsmeur et al., 2018). Due to recent severe rainfall shortages and large-scale sugarcane farming in the sloping field, drought has become one of the most critical constraints of sugarcane growth, development, and productivity. Up to 80% of sugarcane yield is produced during the tillering and grand growth phases which are known to be seriously affected by water scarcity (Gentile et al., 2015). Mining of the genes involved in drought resistance and illuminating the molecular mechanisms that underlie this resistance is of great importance in sugarcane breeding programs. In our study, 26 *SINA* genes were identified from the three genomes of *S. spontaneum, Saccharum spp*. and *Sorghum bicolor*. Their phylogenetic relationship, gene structure, conserved motif composition, chromosomal localization, evolution and cis-regulatory elements were analyzed. Further, we analyzed transcriptome dataset for sugarcane under drought stress and found that a *SINA* gene *SsSINA1a* was significantly induced. Functional characterization indicated that overexpression of *SsSINA1a* increases drought tolerance in Arabidopsis. Hence, this study provides a new candidate gene for sugarcane drought-resistant breeding.

## 2. Materials and methods

### 2.1. Identification and characterization of *SINA* gene family

To identify the members of *SINA* gene family in sugarcane and sorghum, all SINA proteins in Arabidopsis obtained from the TAIR database (https://www.arabidopsis.org/) were used as queries to search sugarcane *S. spontaneum* AP85-441 genome (https://www.life.illinois.edu/ming/downloads/Spontaneum_genome/) (Zhang et al., 2018), the entire *Saccharum* spp. R570 BAC clones (http://sugarcane-genome.cirad.fr/) (Garsmeur et al., 2018) and the *Sorghum bicolor* L. 454_v3.1.1 reference genome (https://phytozome-next.jgi.doe.gov/) (Goodstein et al., 2012) by the BLASTP program with an e-value of 1×e^-5^ as the threshold. A further search was performed with Hidden Markov Model (HMM) analysis using HMMER 3.0 program (Wheeler et al., 2013). The SINA domain (PF03145) HMM profile downloaded from PFAM database (http://pfam.xfam.org/) (Finn et al., 2016) was used to search protein sequences containing SINA domain. The HMMER hits were compared with BLASTP results and parsed by manual editing. Then candidate SINA proteins were submitted to the NCBI Batch CD-search database (http://www.ncbi.nlm.nih.gov/Structure/bwrpsb/bwrpsb.cgi) to confirm the presence and integrity of the SINA domain (PF03145) and the RING finger domain (cd16571). The theoretical isoelectric point (PI), molecular weight (MW), instability index, and grand average of hydropathicity (GRAVY) prediction of the SINA proteins were using the ProtParam tool (http://web.expasy.org/protparam/) (Wilkins et al., 1999). Subcellular localization prediction of the SINA proteins was predicted using the WoLF PSORT web server (https://wolfpsort.hgc.jp/).

### 2.2. Phylogenetic analysis, gene structure and sequence analysis of *SINA* gene family

MEGA-X software was used to perform multiple sequence alignment between the amino acid sequences of SINAs and construct a neighbor-joining evolutionary tree of SINA proteins from 10 plant species with 1000 bootstrap replications (Table S1 and S2). To display the conserved sites of the SINA and RING finger domains of SINA proteins, DNAMAN software was used to visualize multiple sequence alignment result.

The distributions of exons, introns and untranslated regions of each *SINA* gene were obtained from the genome annotation files of *S. spontaneum* AP85-441, sugarcane R570 and *Sorghum bicolor*. The MEME online software (http://meme-suite.org/) (Bailey et al., 2015) was used to search for 1-10 motifs with 6-210 amino acid widths, each containing 5-10 conserved motifs. TBtools was used to visualize gene structure and the MEME result (Chen et al., 2020).

### 2.3. Chromosomal location, collinearity and gene duplication of *SINA* gene family

Chromosomal location information of all *SINA* was obtained from the *S. spontaneum* and *S. bicolor* genome annotation file. Each *SINA* gene mapped to the corresponding chromosome was visualized using TBtools (Chen et al., 2020). Gene duplication events were analyzed using the collinearity and evolutionary analysis tool MCScanX (Wang et al., 2012). The identified collinear gene pairs were mapped to their respective locus in the *S. spontaneum* and *S. bicolor* genome in a circular diagram using Circos 0.69 (Krzywinski et al., 2009). For gene selection pressure analysis, the ratio of non-synonymous (Ka) to synonymous (Ks) substitutions of *SINA* collinearity genes was calculated by TBtools (Chen et al., 2020).

### 2.4. Identification of cis-regulatory elements in *SINA* gene promoters

The 2000-bp promoter sequences upstream of the start codons of all *SINA* genes were extracted and then submitted to the PlantCARE website (http://bioinformatics.psb.ugent.be/webtools/plantcare/html/) (Rombauts et al., 1999) for predicting their regulatory motifs and estimate potentially related functions. TBtools was used to visualize the prediction result (Chen et al., 2020).

### 2.5. Expression profiles of *SINA* gene family

Based on the reported transcriptome data, the *Saccharum officinarum* L. Badila and sugarcane cultivar ROC22 from the National Sugarcane Engineering and Technology Research Center were selected. Drought stress treatment was initiated when healthy single-bud sugarcane stems about 8 cm in length were cultivated to the 3-5 leaf stage. Second leaves from five independent plants were collected at 0, 4, 8, 16 and 32 hours after drought stress treatment and used as samples for RNA-Seq (Li et al., 2022). The *S. spontaneum* genome (Zhang et al., 2018) was used as a reference annotation library for RNA-Seq data analysis. The gene expression level was normalized using the Fragments per Kilobase of transcript per Million mapped reads (FPKM) method. The expression level of *SINA* gene family in sugarcane tissue under drought stress was converted by log2 (FPKM). The heat map of gene expression was generated using TBtools (Chen et al., 2020).

### 2.6. Plant materials and culture conditions

The wild-type (WT) and *SsSINA1a*-overexpressing Arabidopsis lines used in this study were in the Columbia (Col-0) background. Seeds of all Arabidopsis lines were placed in the dark at 4 °C for 2-3 d and then transferred to a growth chambers with a 16/8-h light/dark photoperiod (120 μmol m –2 s–1) at 22–24 °C. Seeds of *Nicotiana benthamiana* were germinated in nutrient soil in a growth chambers with a 16/8-h light/dark photoperiod (120 μmol m–^2^ s–^1^) at 25-28 °C. The seedlings of Arabidopsis and tobacco were subsequently transplanted to new soil plemented with adequate nutrients for further growth under the same conditions.

### 2.7. Subcellular localization analysis of SsSINA1a

The ORF of *SsSINA1a* was amplified and inserted into vector pBWA(V)HS-GFP digested with BsaI and EcoRI to obtain a vector in which green fluorescent protein (GFP) was fused with *SsSINA1a* at its C-terminus, and the 35S::GFP vector was used as a control. The primers used are listed in Table S3. The 35S::SsSINA1a:GFP and 35S::GFP were introduced into *Agrobacterium tumefaciens* GV3101 strains and a suspension with OD600=0.5-0.8 were used to were infiltrated into tabacco leaves. After 48-60 h. the infiltrated leaves were observed by laser confocal scanning micro scope (LCSM).

### 2.8. Plant transformation of *SsSINA1a* and drought assay

The *Agrobacterium tumefaciens* (strain GV3101) carried 35S::SsSINA1a:GFP was used to transform Arabidopsis according to previously described method (Hao et al., 2012).The transgenic T0 Arabidopsis seeds were first sown on 1/2MS medium, and the normally growing T1 plants were transplanted into nutrient soil to continue to grow. The T1 generation seeds were cultured on the screening medium, and the T2 lines statistically consistented with the 3:1 segregation ratio were screened. Total RNA of 3-week-old *SsSINA1a*-overexpressing Arabidopsis was extracted using an RNA Extraction Kit (DP411, Tiangen Biotech). The housekeeping genes *AtACTIN* was used as the internal control. The primers used are listed in Table S3. The 3-week-old WT and *SsSINA1a*-overexpressing Arabidopsis seedlings grown in nutrient soil were stopped watering for 3-4 weeks, then rewatered for 2-3 days, finally the survival rate was calculated. The rosette leaves of 4-week-old WT and *SsSINA1a*-overexpressing Arabidopsis were separated and placed in uncovered petri dishes under dim light and room temperature. Then, the weight of each rosette leaf was measured at different time points (0, 2, 4 and 6 h).

## 3. Results

### 3.1. Identification and characterization of *SINA* gene family

Using five SINA proteins of Arabidopsis (AtSINA1-AtSINA5) as the query sequences, a total of 26 *SINA* genes were identified, including 15 *SsSINAs* in *S. spontaneum*, 5 *ShSINAs* in *Saccharum spp*. R570, and 6 *SbSINAs* in *S. bicolor* (Table S4). By comparing phylogenetic relationships and protein identity with sorghum homologs and paralogs, *S. spontaneum* representative gene models for different alleles have been screened now (Zhang et al., 2018). Tandem duplicated genes and paralogs are considered novel genes, and their gene IDs are followed by T and P, respectively (Yuan et al., 2021). All identified *SINA* genes are named according to the physical position on the chromosomes or BACs. The alleles of *SsSINAs* were marked the letters “a,” “b,” “c,” and “d” (Table S4). Among these 15 *SsSINA* genes (*SsSINA1-10*), the *SsSINA1* gene had three alleles (*SsSINA1a/1b/1c*) and *SsSINA2/4/6* genes had two alleles. However, other *SINA* family members of *S. spontaneum* have no alleles, suggesting that alleles may be lost in the process of evolution. Analysis of the primary structure of protein shows that, the protein lengths of SsSINA1–10, ShSIAN1–5, and SbSINA1–6 were 301–348, 302–351 and 302–353, respectively (Table S4). Among the 26 SINA proteins, 16 SINA proteins were predicted to be acidic and 13 SINA proteins were predicted to be stable or unstable (Table S4). The grand average of hydropathicity (GRAVY) of all SINA proteins were all less than 0 (Table S4). In addition, the prediction of subcellular localization showed that 26 SINA proteins were located in the nucleus, cytoplasmic or chloroplast (Table S4).

### 3.2. Phylogenetic analysis of the *SINA* gene family

We performed a phylogenetic analysis with 76 SINA proteins from *S. spontaneum, Saccharum* spp. R570, *S. bicolor, A. thaliana*, rice (*Oryza sativa*), maize (*Zea mays*), tomato (*Solanum lycopersicum*), poplar (*Populus trichocarpa*), apple (*Malus domestica*) and *Medicago truncatula* to gain additional insight into the evolutionary relationships of *SINA* gene family (Table S1 and S2). The phylogenetic tree constructed by the neighbor-joining analysis divided all SINA proteins into three groups (Fig.1). Our 26 SINAs were clustered into three groups as follows: 11 within group I; 11 within group □; 4 within group □. The SINA proteins of sugarcane (SsSINAs and ShSINAs) have a closer evolutionary distance to that of sorghum and maize than rice and Arabidopsis, and have the closest genetic distance to sorghum.

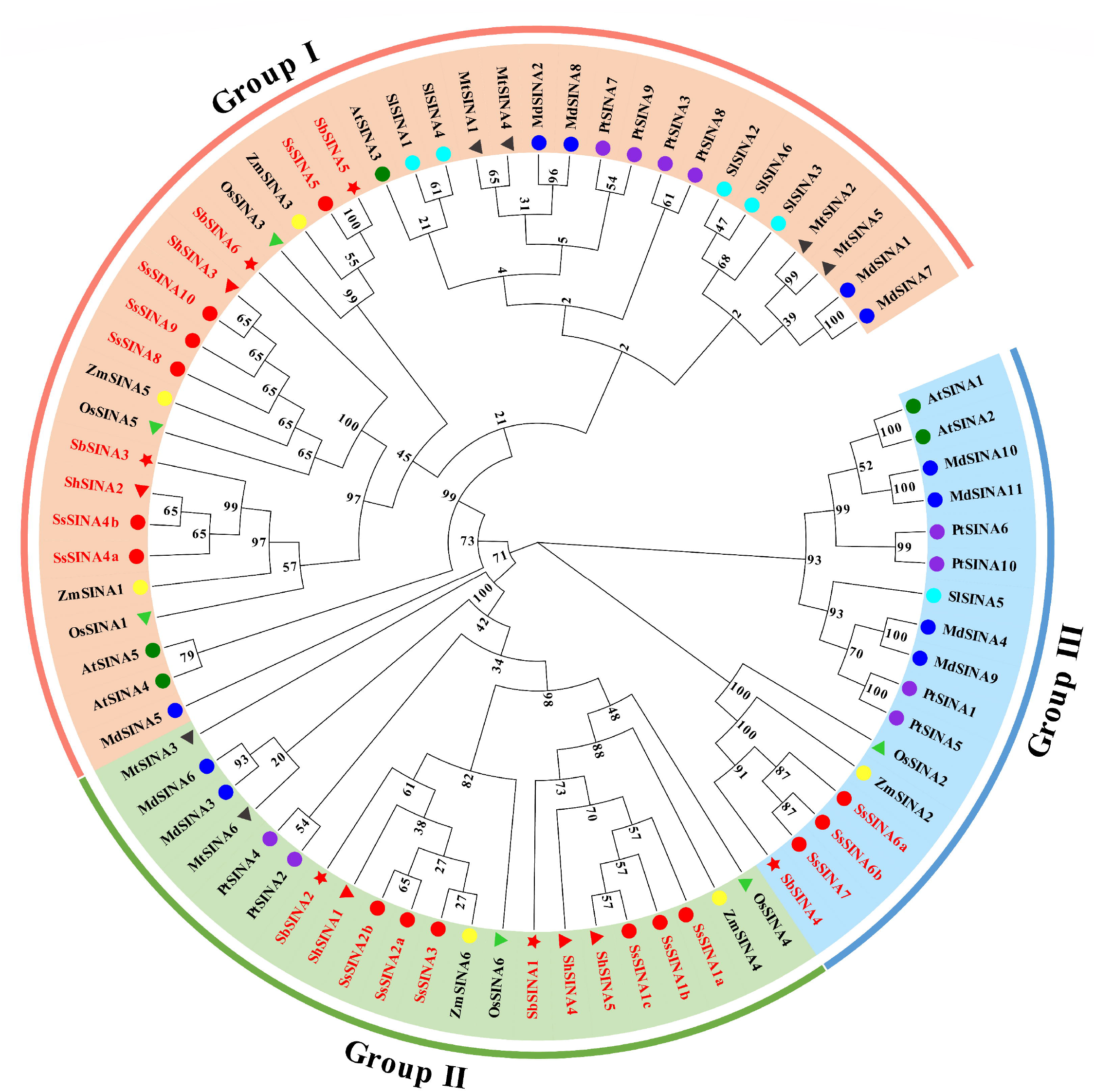

### 3.3. Gene structure and conserved motifs of the *SINA* gene family

The exon-intron structure of the 26 *SINA* genes was analyzed to provide further insight into the evolution of these genes. The results revealed that all *SINA* genes have different number of exons/introns, in which 18 *SINA* (account for 69.2%) possess three exons and two introns, and other 8 *SINA* (account for 30.8%) possess four exons and three introns (Fig. 2). Additionally, although the protein sequence identity between *SsSINA1a, SsSINA1b* and *ShSINA5* is 100%, the structures of their related gene were different (Table S10), suggesting that the three genes may have different functions in the process of evolution.

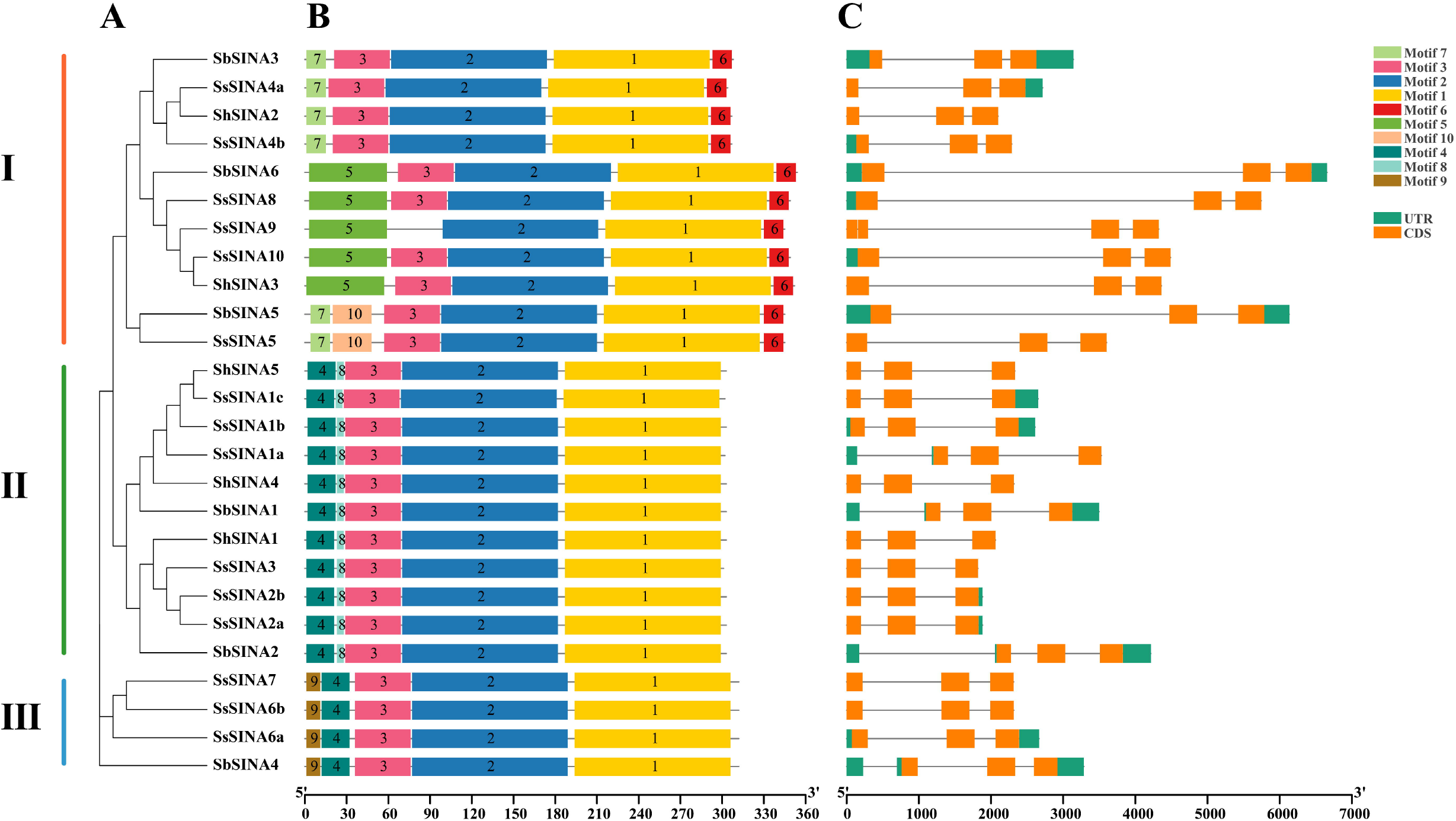

The conserved motifs of 26 SINA proteins were confirmed by MEME, and 10 distinct motifs were identified (Fig. 2 and Table S5). All 26 SINA proteins contained conserved motifs 1 and 2; except for SsSINA9, conserved motif 3 was also ubiquitous in the SINA proteins (Fig. 2). Meanwhile, the motif distribution of SINA proteins in the same branch of the phylogenetic tree is highly similar (Fig. 2). At the proteins C terminus, all group I members have motif 6, while all group II and group □ members lack this motif (Fig. 2). Both group II and group □ proteins have motif4 at the N terminus and motifs 8 and motif 9 were the unique motifs in group II and group □, respectively (Fig. 2). The multiple sequence alignment found that all SINA proteins have conserved cysteine-rich C3HC4-type (Cys-X2-Cys-X-Cys-XHis-X-Cys-X2-Cys-X-Cys-X2-Cys) RING finger domain and SINA domain, while the N-terminal region is highly variable (Fig. S1). The similar distribution of motifs suggests that the *SINA* genes are relatively conserved during evolution. And the uniqueness of the motif distribution in SINA proteins of different group also reflects the differentiation of *SINA* genes function during evolution.

### 3.4. Chromosomal location, collinearity and gene duplication of *SINA* gene family

Due to the large number of unassembled BAC clones in the R570 genome, we only analyzed the chromosomal location and gene replication type of *SINA* gene family in *S. spontaneum* and *S. bicolor*. 15 *SsSINAs* are distributed on 12 chromosomes of *S. spontaneum* in which chromosome 2A (Ss2A), Ss4D and Ss7D each contained two *SsSINAs*; the remaining 9 chromosomes each contained one *SsSINA* gene, and there are no *SINA* genes in the three homologous chromosome groups of Ss5, Ss6 and Ss8 (Fig. S2). The 6 *SbSINAs* were unevenly distributed on five chromosomes of *S. bicolor*. Two *SbSINAs* (*SbSINA4/5*) were located on *S. bicolor* chromosome 4 (Sb04). The remaining four *S. bicolor* chromosomes each had one *SbSINA* gene. In 15 *SsSINAs*, 12 whole-genome or segmental, 1 proximal, 1 dispersed, and 1 tandem duplication genes were detected. In six *SbSINAs*, five whole-genome or segmental and one dispersed duplication genes were detected (Table S6).

Meanwhile, the Collinearity relationships of the *SINA* gene family in *S. spontaneum* and *S. bicolor* was analyzed (Fig. S3). A total of 11 collinearity gene pairs in 15 *SsSINAs* were found, of which 5 pairs (45.45%) occurred between alleles, and 6 pairs (54.55%) occurred between non-alleles (Fig. S3A and Table S7). And 4 collinearity gene pairs in six *SbSINAs* were also found. There were 11 orthologous *SINA* gene pairs between *S. spontaneum* and *S. bicolor*. The Ka/Ks ratio in *SINA* collinearity gene pairs ranged from 0.00 to 0.44. All collinearity gene pairs had a Ka/Ks ratio < 1, suggesting that *SINA* gene family may have experienced quite strong pressure of purifying selection during evolution.

### 3.5. Identification of cis regulatory elements

In the 2000-bp promoters of the 26 *SINA* genes, four types of cis-regulatory elements were predicted, including phytohormone-, plant growth/development-, light- and stress-related elements (Fig. 3, Table S8 and S9). The ABA-responsive elements existed in the promoters of 88.46% *SINA* genes. In addition, the promoters of 22 (84.61%), 14 (53.84%), 10 (38.46%) and 9 (34.62%) *SINA* genes contained MeJA-, gibberellin (GA)-, auxin-, and salicylic acid (SA)-responsive elements, respectively. Low-temperature elements (LTRs), defense and stress responsive elements (TC-rich repeats) and MYB binding sites (MBSs) involved in drought-inducibility were found in the promoters of 18 (69.23%), 11 (42.31%) and 9 (34.62%) *SINA* genes, respectively. For plant growth and development, promoters of 8 (30.77%) *SINA* genes contained CAT-boxes elements which is an element regulating meristem expression. Promoters of *SsSINA4b* and *ShSINA2* may have been involved in circadian control. These results indicate that the *SINA* gene family may be widely involved in the response to various stress and in the regulation of plant growth and development.

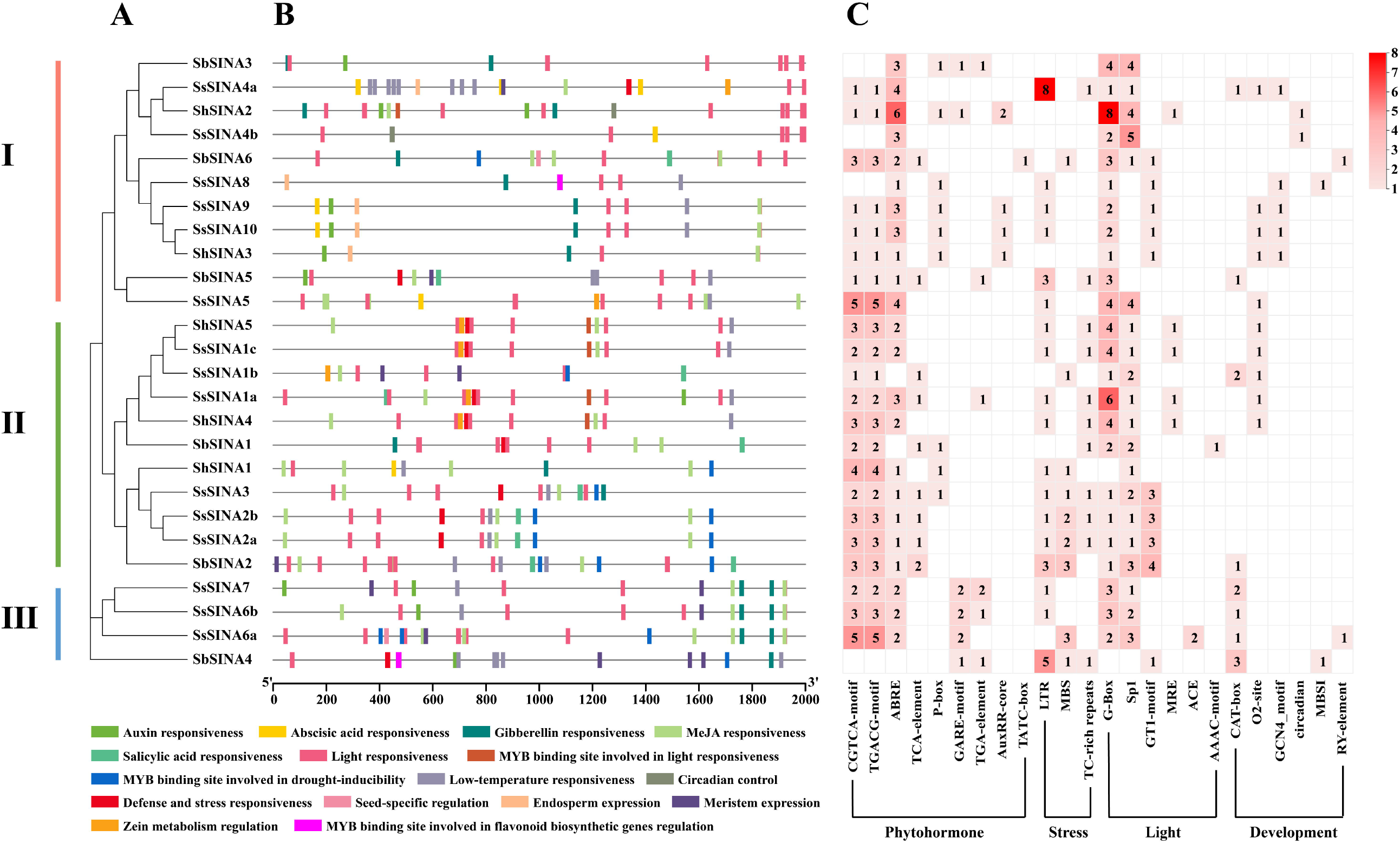

### 3.6. Expression profiles of *SINA* genes in sugarcane under drought stress

To identify important candidate *SINA* genes in sugarcane response to abotic stress, we analyzed reported RNA-seq dataset derive from sugarcane cultivar Badila and ROC22 with drought stress treatment (Li et al., 2022). The results show that the expression level of *SsSINA* genes in group □ was generally lower than that in group □, and the *SsSINA2a/2b/3* in group □ were almost not expressed in two sugarcanes (Fig. 4). Additionally, we found that the expression patterns of *SsSINA1a/1b/1c* were similar under drought stress, and their expression levels were all significantly up-regulated (fold change > 2), suggesting that these three group □ members may play an important role in sugarcane response to drought. Subsequently, *SINA1a* with the highest expression was selected to further study its biofunction in drought stress.

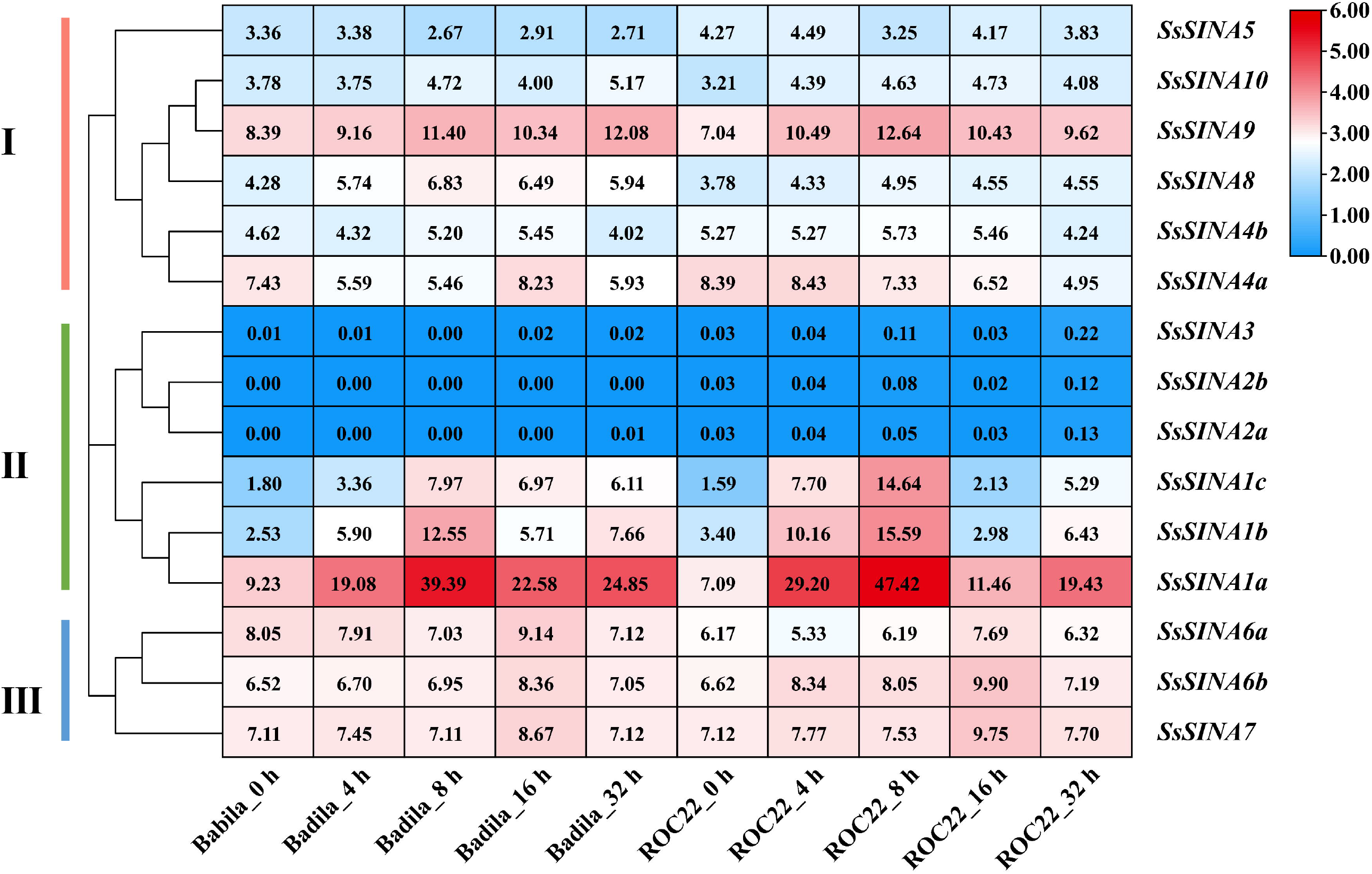

### 3.7. Subcellular localization analysis of SsSINA1a

To verify the prediction results of SsSINA1a protein subcellular localization, the full-length of *SsSINA1a* was inserted into the pBWA(V)HS-GFP vector in which *SsSINA1a* fused with GFP protein driven by the 35S promoter, and then expressed in *N. benthamiana* leaves. Subcellular localization results showed that the GFP signal of the empty pBWA(V)HS-GFP vector is expressed in various organelles in *N. benthamiana*, and the GFP signal of the recombinant vector 35S::SsSINA1a:GFP is only generated in the nucleus (Fig. 5), suggesting that SsSINA1a is a nuclear protein which was consistent with our prediction.

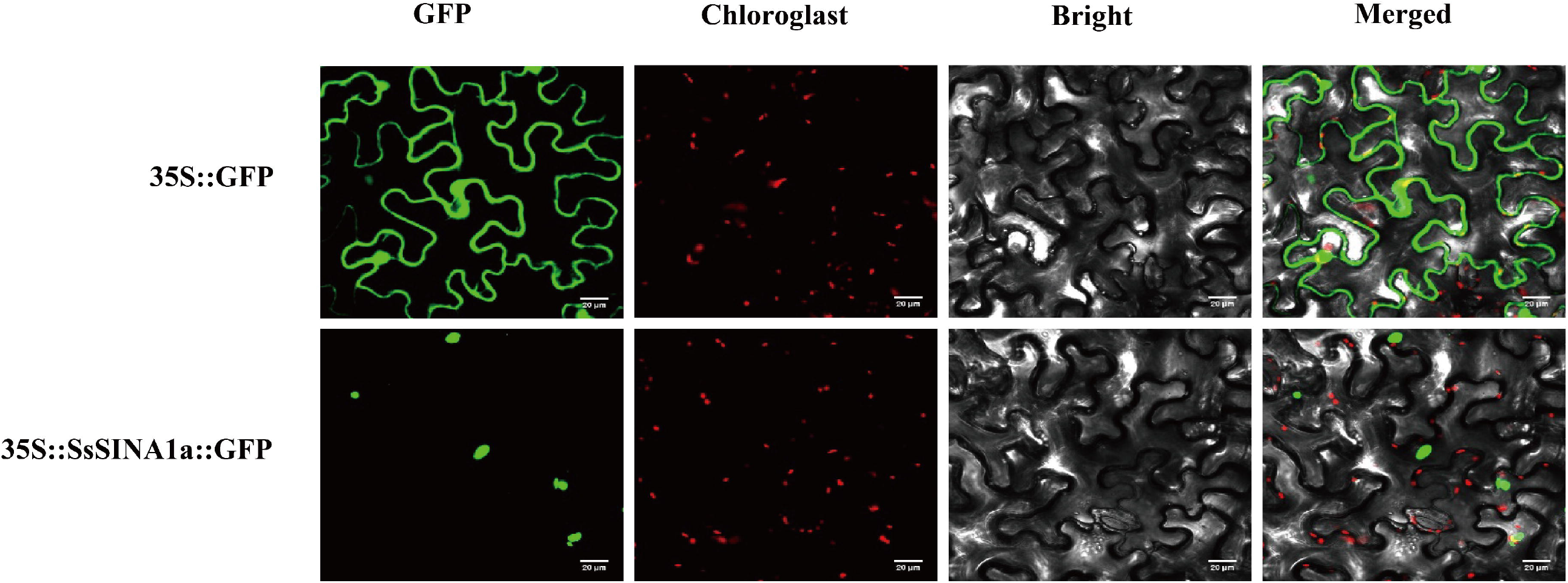

### 3.8. SsSINA1a increases drought tolerance in Arabidopsis

To study the biofunction of *SsSINA1a* in response to drought stress, we performed heterologous overexpression of *SsSINA1a* under the 35S promoter in Arabidopsis, and two independent transgenic lines (*SsSINA1a*-*OE4* and *SsSINA1a*-*OE10*) with high expression levels were selected (Fig. 6A). After 28 days of ceasing watering, the wilting degree of WT plants was higher than that of *SsSINAT1a-OE4* and *SsSINAT1a-OE10* lines (Fig. 6B). After 3 days of rehydration, the majority of WT seedlings have stopped growing even death, while most of *SsSINAT1a-*overexpressing seedlings recovered normal growth (Fig. 6B). Further statistics showed that the survival rate of WT Arabidopsis was only 6.7%, while that of the transgenic lines was 61.1% (Fig. 6C). Moreover, water content of the detached leaf in *SsSINA1a-OE4* and *SsSINA1a-OE10* was significantly higher than WT (Fig. 6D). These results suggested that the overexpression of *SsSINA1a* improved the drought resistance of Arabidopsis.

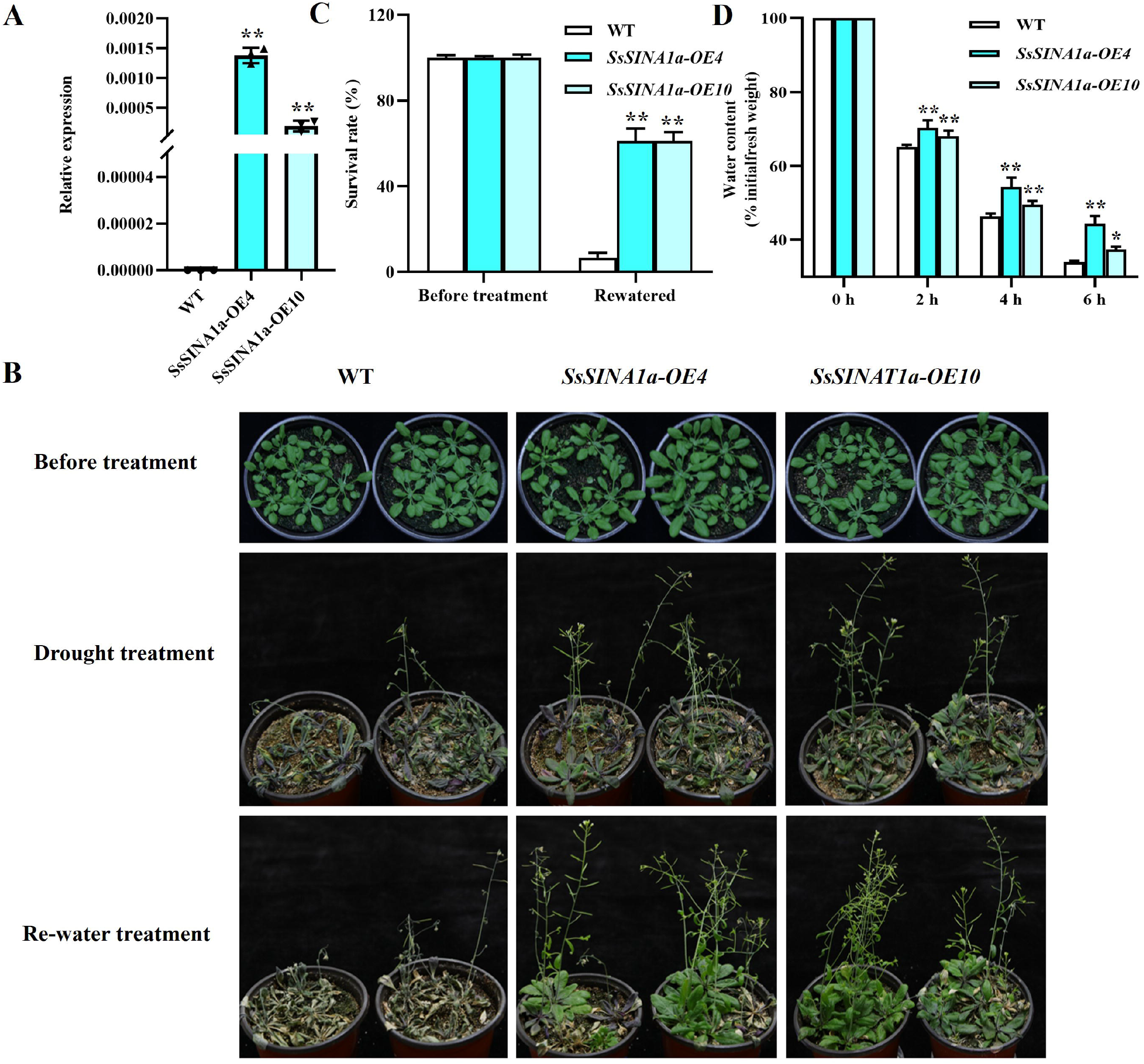

## 4. Discussion

As an important E3 ubiquitin ligase, SINA and its homologues proteins can recognize a variety of different substrate proteins and make them ubiquitinated and degraded (Qi et al., 2013; Chen et al., 2013). The function of SIAH proteins in animals and humans has been widely studied, especially in the field of medicine, and these studies revealed that SIAH proteins are the potential therapeutic targets for various diseases including cancer (Roperch et al., 1999; Liu et al., 2008; Cai et al., 2015; Siswanto et al., 2018). In recent years, increasing studies reported the function of SINA proteins in regulation of plant growth and the response to different stresses in plants (Zhang et al., 2019). In this study, the characteristics of the *SINA* gene family in sugarcane and regulatory role of *SsSINA1a* in response to drought stress were analyzed.

The 26 SINA proteins identified in this study all contain highly conserved RING finger domain and SINA domain (Fig. S1). The evolutionary relationship of SINA proteins divided the of sugarcane *SINA* gene family into three groups (Fig. 1). Group I contains the largest number of members with slightly long C□terminus and motif 6 is a unique conserved motif in group I, which is similar to the results of previous studies (Fig. 2B) (Wang et al., 2008; Wang et al., 2018; Li et al., 2020). There is a small difference in the motif between the group □ and group □ at the N-terminal, suggesting that functional differences may occur in the process of evolution. In addition, we found that the SINA proteins of monocotyledonous and dicotyledonous plants were clustered separately in phylogenetic tree, indicating that the *SINA* gene family originated later than the differentiation of monocotyledonous and dicotyledonous plants (Fig. 1). Same as the results of previous studies, the second CDS of our most *SINA* genes (96.15%) show quite strong length conservation (387bp), which also confirms that some functional domains of *SINA* are very conservative in the process of evolution (Wang et al., 2008). Gene replication is a remarkable feature of plant genome structure and the main mechanism driving the expansion and evolution of gene family (Flagel et al., 2009; Wang et al., 2018). In Arabidopsis, poplar and apple, the expansion mechanism of *SINA* gene family is mainly segmental replication, which is more obvious in the polyploid sugarcane (Table. S6) (Wang et al., 2008; Li et al., 2020). Five duplicated gene pairs among *SsSINA* alleles showing similar gene structures and motif patterns that imply some degree of functional redundancy, which is verified in the expression pattern from the transcriptome (Table. S7 and Fig. 4).

All promoters of 26 *SINA* genes contains plant hormone response elements, plant growth and development related elements and stress response elements, which mean the *SINA* gene family is widely involved in the regulation of various physiological processes (Fig. 3). There are some differences in the type and number of cis-elements among members of the *SINA* gene family, indicating that gene replication events produce new functions on the basis of retaining the original functions in the process of evolution, which makes *SINA* have a wider range of biological activities (Table. S9). As an important member of ubiquitin-proteasome pathway, SINA E3 ligase has been proved to have extensive and rich regulatory effects in many plants, including the growth and development of plant tissues (Xie et al., 2002; Park et al., 2007; Park et al., 2010; Wang et al., 2018), secondary metabolism (Welsch et al., 2007), symbiotic nitrogen fixation (Den et al., 2008), autophagy and drought stress responses (Ning et al., 2011; Bao et al., 2014; Qi et al., 2017), cold stress responses (Fan et al., 2017), plant immunity (Miao et al., 2016; Wang et al., 2018; Ren et al., 2021) and so on. Plant SINA proteins have been studied to be located in the nucleus or cytoplasm in the past (Wang et al., 2018; Li et al., 2020; Xia et al., 2020; Ren et al., 2021). In this study, we identified a significantly drought-induced gene *SsSINA1a* located in the nucleus (Fig. 5). And the overexpression of *SsSINA1a* reduced the water loss rate of leaves and improved the drought tolerance of Arabidopsis (Fig. 6).

Although many members of the *SINA* gene family have been identified so far, there are still few studies on the function of *SINA* response to drought stress, and its regulatory mechanism is elusive. Here, we identified a sugarcane RING E3 ligase SsSINA1a that plays a key role in regulating the drought response of sugarcane and Arabidopsis. And this study advances the functional study of the sugarcane *SINA* gene family and provides a new insight for the target genes research of sugarcane drought-resistant breeding.

## Acknowledgments

The authors thank all editors and reviewers for their comments on this manuscript.

