## Supplementary figures and images for "Genome-wide identification of *SINA* gene family in sugarcane and functional analysis of *SsSINA1a* in drought response"

### Supplemental Fig. S1

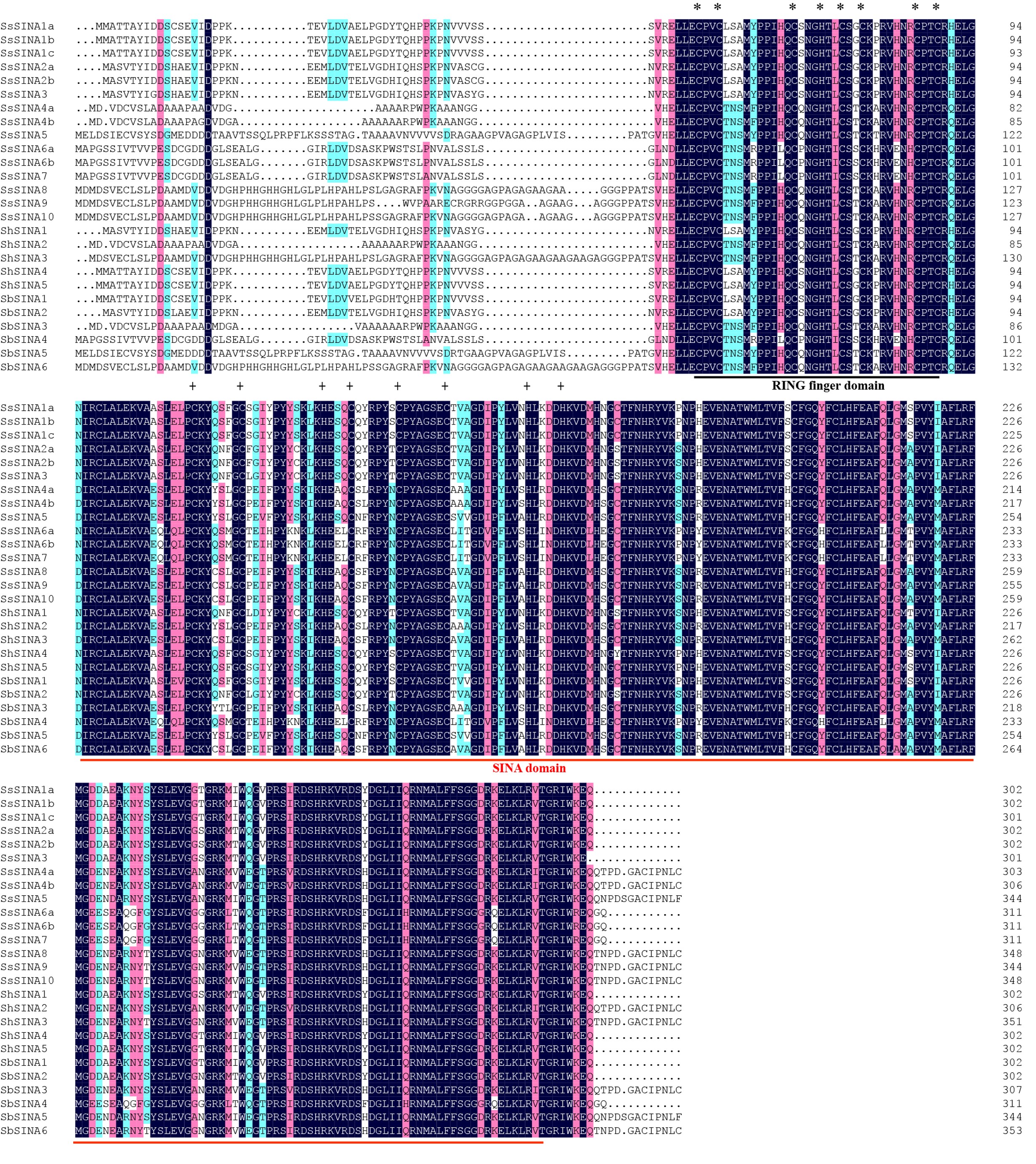

### Supplemental Fig. S2

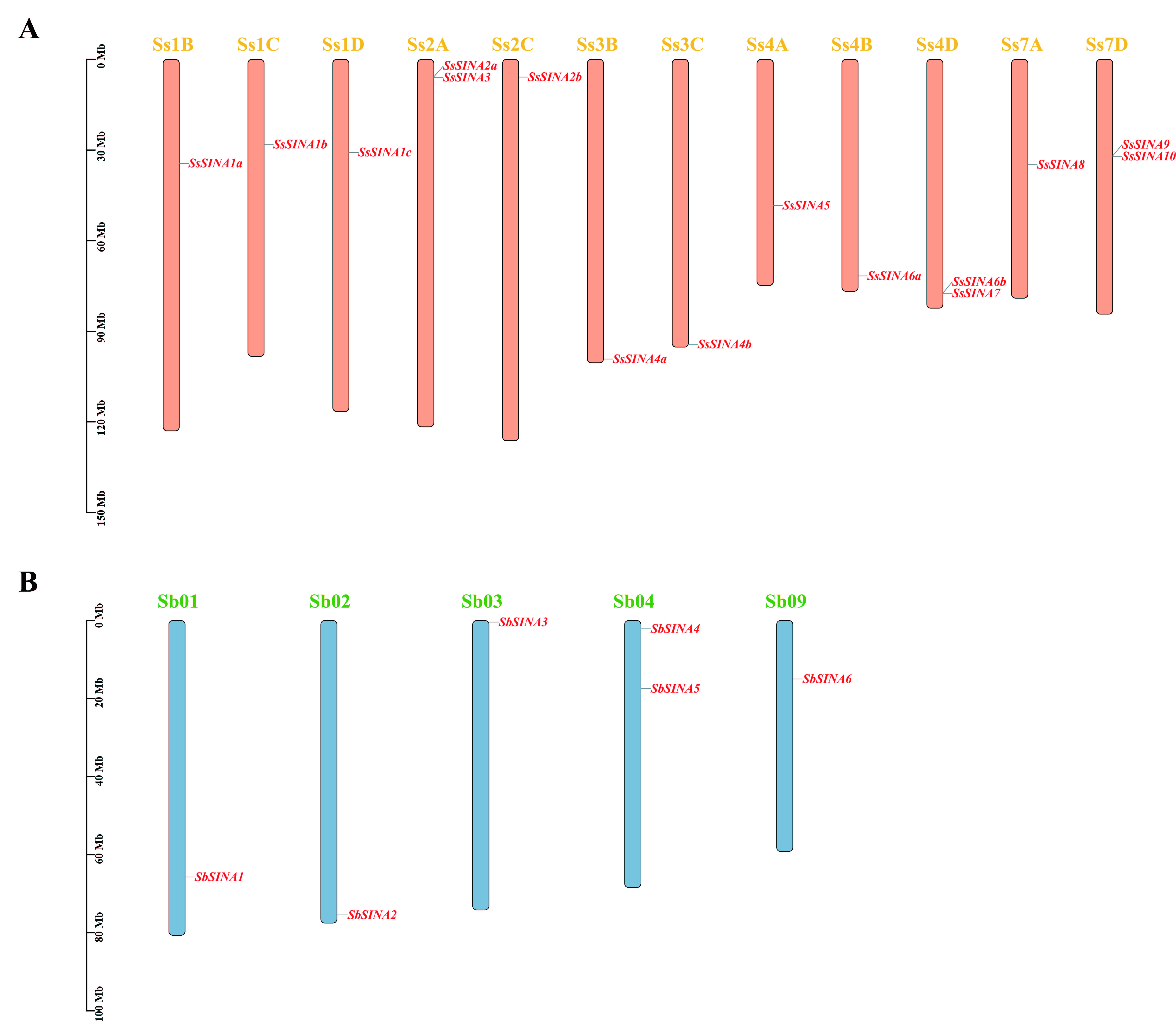

### Supplemental Fig. S3

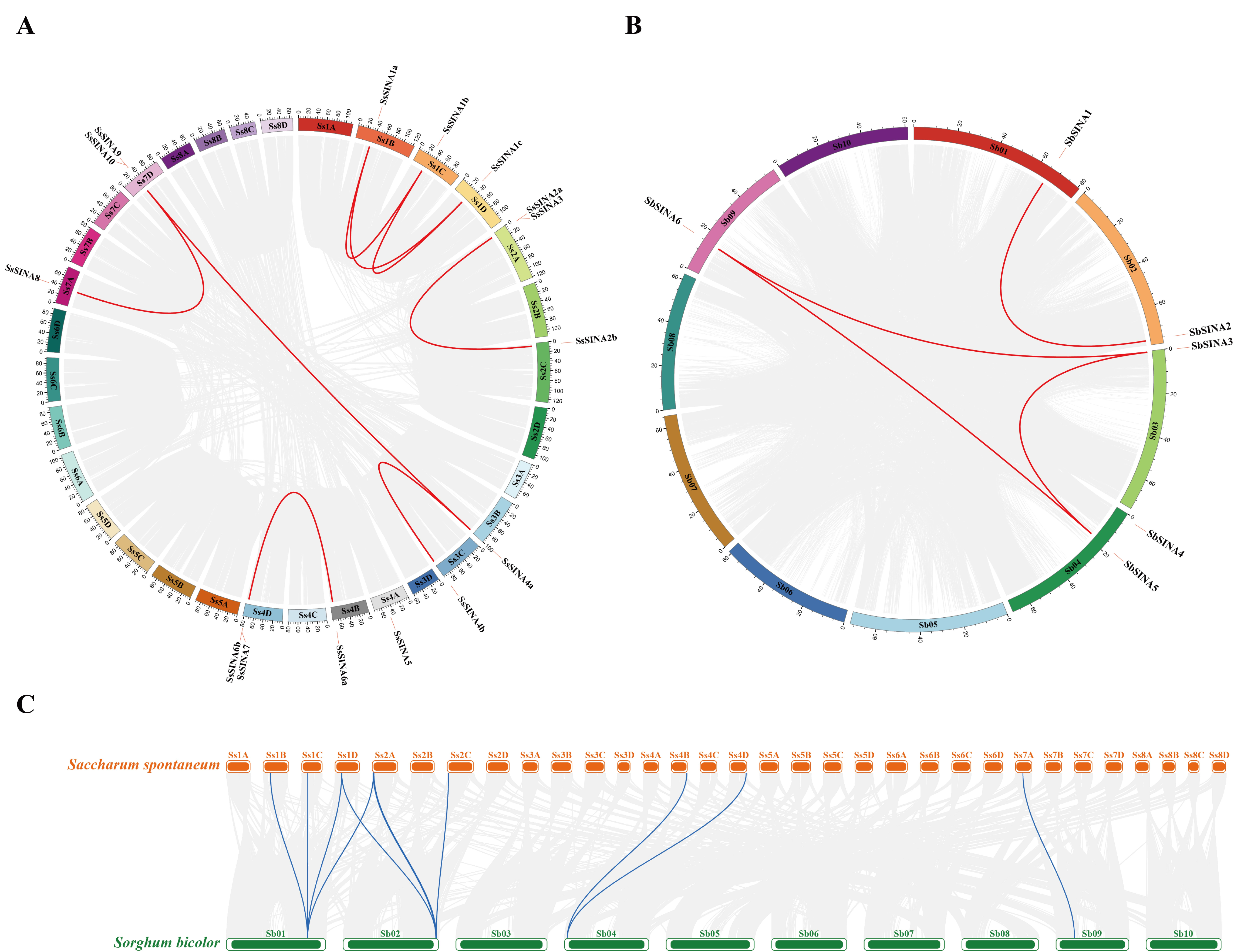
